# MicroRNA840 accelerates leaf senescence by targeting the overlapping 3’UTRs of *PPR* and *WHIRLY3* in *Arabidopsis thaliana*

**DOI:** 10.1101/2020.10.23.353052

**Authors:** Ren Yujun, Wang Wanzhen, Lan Wei, Schenke Dirk, Cai Daguang, Miao Ying

## Abstract

MicroRNAs (miRNAs) negatively regulate gene expression by cleaving the target mRNA and/or impairing its translation, thereby playing a crucial role in plant development and environmental stress responses. In Arabidopsis, *MIR840* is located within the overlapping 3’UTR of *PPR* and *WHIRLY3 (WHY3),* both being predicted targets of miR840. Gain- and loss-of-function of miR840 in Arabidopsis resulted in opposite senescent phenotypes. Highest expression of *pri-miR840* is observed at senescence initiation, and is negatively correlated with a significant reduction of *PPR* transcripts but not of *WHY3.* Although *WHY3* transcript levels were not significantly affected by miR840 overexpression, its protein synthesis was strongly reduced. Mutating the cleavage sites or replacing the target sequences abolishes the miR840-mediated degradation of *PPR* transcripts and inhibition of *WHY3* translation. In support for this, concurrent knock-down of both *PPR* and *WHY3* in the WT resulted in the senescent phenotype resembling that of the miR840-overexpressing mutant. This indicates that both PRR and WHY3 are targets in the miR840-regulated senescent pathway. Moreover, single knockout mutant of *PPR* or *WHY3* shows a convergent up-regulated subset of senescence-associated genes, which are also found among those induced by miR840 overexpression. Our data provide evidences for a regulatory role of miR840 in plant senescence.

**Highlight:** MicroRNA840 (miR840) has a unique miRNA-target configuration regulating *PPR* and *WHIRLY3* genes in Arabidopsis. MiR840 is highly expressed at the onset of plant senescent stage. Both *PPR* and *WHIRLY3* transcripts are specifically targeted *in vivo* within their 3’UTR region by mature miR840 or its star strand *in vivo.* Interestingly, *PPR* expression is mainly repressed on mRNA transcript level by cleavage, while WHIRLY3 is predominantly translationally inhibited. We conclude that miR840 enhances plant senescence *via* post transcriptional gene silencing of *PPR* and *WHIRLY3*, which appear to be novel negative joint regulators of plant senescence.

Footnote: The author(s) responsible for distribution of materials integral to the findings presented in this article in accordance with the policy described in the intructions for Authors is: Ying Miao

## Introduction

Senescence is the last developmental stage of whole plants or their organs, and is often associated with a transition, which can be also stimulated by environmental stress. In monocarpic crops, premature senescence leads to the reduction of product yield and postharvest quality. During plant senescence, genes coding for proteins related to autophagy, chlorophyll and lipid catabolism, carbohydrate and nitrogen transport, as well as those involved in generation of reactive oxygen species are up-regulated, whereas others related to protein synthesis and maintenance of mitochondrial or chloroplast functions such as light harvesting, carbon fixation and photorespiration are down-regulated (Lim et al., 2003; Guo and Gan, 2014). Global transcriptome analyses in Arabidopsis showed that approximate 12-16% genes are regulated differentially in senescence-related physiological and pathological processes, indicating the occurrence of extensive transcriptional reprogramming during plant senescence (Guo et al., 2004; Zentgraf et al., 2004; Buchanan-Wollaston et al., 2005; Breeze et al., 2011). This involves tight control by a number of transcription factors (TFs), epigenetic modifications and small non-coding RNAs (Schippers, 2015; Kim et al., 2016; Ren and Miao, 2018; Woo et al., 2019b).

MicroRNAs (miRNAs) are a class of highly conserved endogenous small non-coding RNAs (usually 20-24 nt). Since identification of the first miRNA, *lin-4,* in *Caenorhabditis elegans* (Lee et al., 1993), thousands of miRNAs have been identified in animals and plants, and showed manifold roles in controlling diverse biological processes (Ameres and Zamore, 2013; Dexheimer and Cochella, 2020). In plants, multiple factors contribute to the biogenesis, conversion, mobilization and action mechanisms of miRNAs, and in turn, miRNAs control cognate target genes through transcript cleavage and translational repression (Rogers and Chen, 2013; Xie et al., 2015; Yu et al., 2017). Fewer miRNAs with particular link to the regulation of plant senescence have been functionally charasterized (Woo et al., 2019a). One example is miR164 which targets the NAC domain-containing proteins such as ORE1 and NAC1 to regulated leaf senescence and cell death during development (Kim et al., 2009; Li et al., 2013). Another microRNA, miR319, negatively controls a set of *TCP (TEOSINTE BRANCHED/CYCLOIDEA/PCF)* transcription factor genes, which regulate biosynthesis of the hormone jasmonic acid, to affect leaf development and senescence progression (Schommer et al., 2008). Recently, by using high-throughput smallRNA sequencing strategies, a number of senescence inducible miRNAs in rice, maize and Arabidopsis plants are also discovered (Xu et al., 2014; Thatcher et al., 2015; Qin et al., 2016; Wu et al., 2016). These researches provide large data sets of miRNAs associated with developmental and senescent stages, and in response to nutrition availability or stress conditions. However, specific role of miRNA in controlling senescence of an organ is rarly reported.

Among the senescence-associated miRNAs in leaves (Xu et al., 2014), miR840, is firstly identified in a previous high-throughput pyrosequencing (Rajagopalan et al., 2006), which appears only in genomes in cruciferous plants of the genus Arabidopsis thus considering as an evolutionary young microRNA. A canonical candidate target gene of miR840 is a *WHIRLY3 (WHY3),* which is a less-studied member of the three-gene family of single-stranded-DNA-binding proteins in Arabidopsis (Cappadocia et al., 2013), The *WHIRLY* family includes the well-known leaf senescence regulator *WHIRLY1 (WHY1)* (Miao et al., 2013), the closest paralog of *WHY3.* However, the function of miR840 is still unclear.

The locus *Ath-miR840 (At2g02741)* is located within the 3’UTR region of the protein-coding gene *PPR (At2g02750),* overlapping with the distal portion of the 3’UTR from the opposite strand-encoded gene *WHY3 (At2g02740),* both being predicted targets of miR840 (Rajagopalan et al., 2006). This special locus arrangement categorizes *miR840* into the G3A group of *miRNAs,* which qualitative and quantitative analysis by sequencing are often hindered by the overlapping or adjacent gene transcripts (Armenta-Medina et al., 2017). Here, we demonstrate that miR840 regulates the onset of plant senescence via targeting *PPR* and *WHY3* in two different manners by degradating *PPR* transcripts and inhibiting the *WHY3* translation concurrently. Neither PPR and WHY3 have been implicated in plant senescence so far, but our analysis suggests that they might act in concert to negatively regulate this process since the *WHY3* and *PPR* double mutant *(kdwhy3 appr)* resembles the early senescence phenotype observed in the miR840 overexpression mutant.

## Results

### MiR840 is processed by three Dicer-like ribonucleases (DCLs) with various efficiency in Arabidopsis

The Arabidopsis miRNA840 precursor *(pre-miR840)* gene was predicted to be located within a *PPR* and *WHY3* cross-locus (Figure 1A-B), and belonged to group G3A miRNAs (Rajagopalan et al., 2006; Lepe-Soltero et al., 2017). The abundance of the mature miR840 was reported to be reduced by about 0.9-fold in the *dcl1* mutant embryos (Nodine and Bartel, 2010; Armenta-Medina et al., 2017). Accordantly, mature miR840 was hardly detected by RNA gel blot analysis in rosettes of the *dcl1* mutant, as compared with the small interfering RNA *siRNA1003* serving as a control, which is known not to be affected by *dcl1* mutation (Figure 1C, left panel). To test whether the other three Arabidopsis *DCL* genes are involved in the production of miR840, we quantified mature miR840 levels additionally in the *dcl2, dcl3, dcl4, dcl4-2t* single and *dcl2 dcl4 (dcl2&4)* double mutants (Pelissier et al., 2011) using stem-loop semi- and qRT-PCR. Except for the *dcl3*, in which the miR840 level was comparable to that in WT plants, *dcl2* and *dcl4* showed a decrease in miR840 abundance approximately 39% and 73% relative to the WT, respectively. The double mutant *dcl2/dcl4* exhibted a reduction of miR840 levels by 83% as compared to the WT (Figure 1C-D). As controls, the expression of the known DCL1-dependent *miR173* was unaffected in the *dcl2, dcl3, dcl4* single and *dcl2 dcl4* double mutants, whereas the expression of the DCL4-dependent *miR839* was strongly declined in *dcl4* (by 95%) and in *dcl2 dcl4* (by 97%), as well as in the *dcl2* single mutant (by 61%) (Figure 1C-D) in consistence with a previous study (Pelissier et al., 2011). Therefore, we conclude that the production of the mature miR840 is dependent mainly on DCL1 and to a lesser extend also on DCL2 and DCL4

**Figure 1.**
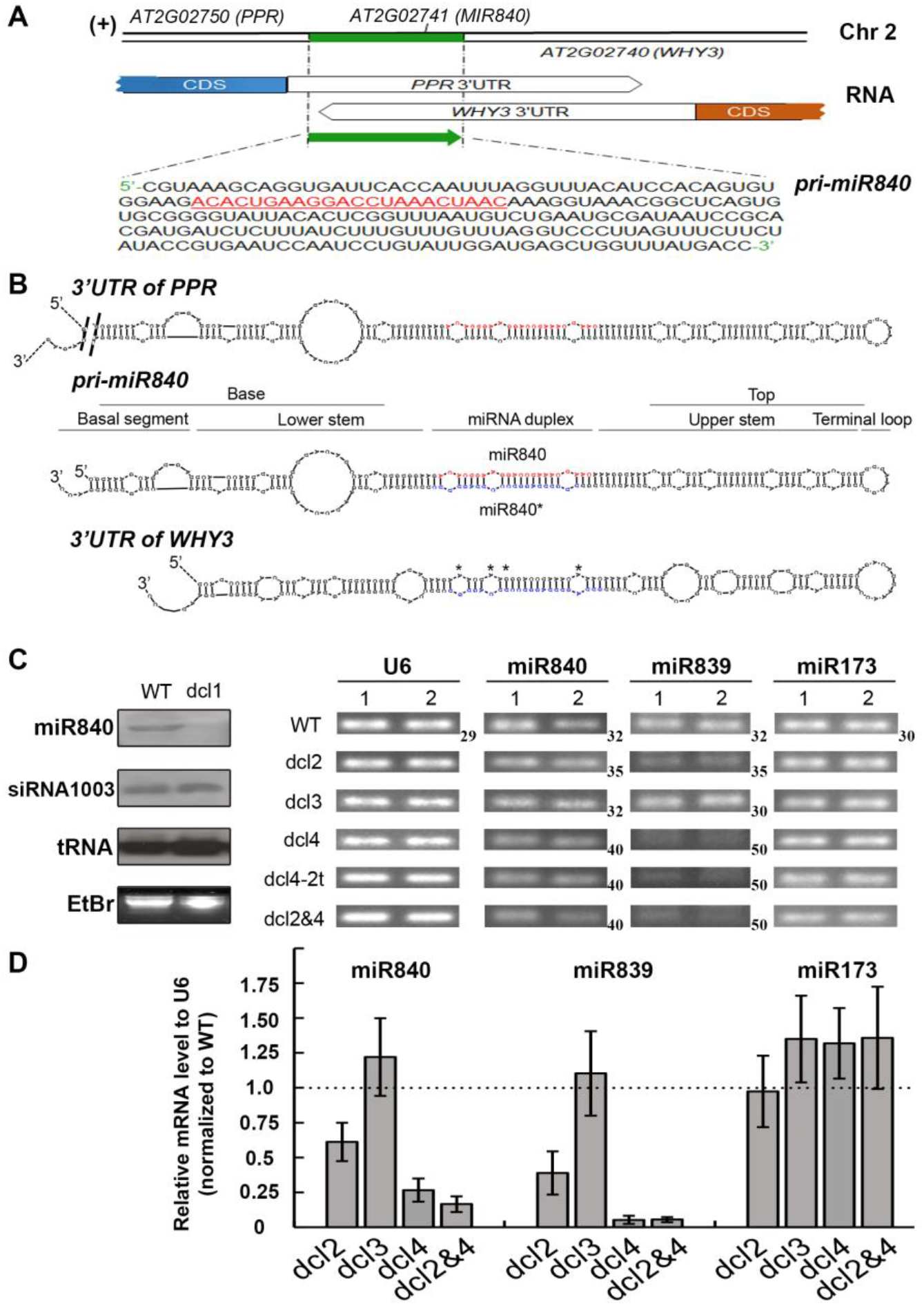
Gene locus and DCL-dependency of miR840 in Arabidopsis. **(A)** The *MIR840* is located within a convergent gene pair in chromosome 2. The 226 bp *pri-miR840* was shown and mature miR840 sequence was underlined and highlighted. **(B)** Predicted RNA structure of the *PPR 3’UTR, pri-miR840* and *WHY3 3’UTR,* by RNAstructure© ver.6.0.1. The mature miR840 strand (red) and its pairing strand *miR840\** (blue) were labeled. **(C)** Northern blot (leaf panel) and stem-loop semi-qPCR (right panel) detection of miR840 in *dcl1*, *dcl2*, *dcl3*, *dcl4*, *dcl4-2t* and double *dcl2dcl4* mutant plants. EtBr: ethidium bromide staining. For semi-qPCR, the PCR cycles for amplification are shown beside the gel of each microRNA with unequal numbers indicated. **(D)** Stem-loop RT-qPCR showing fold-change of miR840 level in *dcl2, dcl3, dcl4* and double *dcl2dcl4* mutant plants over the that in WT.

### *MIR840* expression in rosettes reaches its maximum at the onset of plant senescence

To identify the role of miR840 in regulating plant senescence, we first analyzed the tissue-specific expression of the miR840 precursor (*pri*-*miR840*) together with the target genes *PPR* and *WHY3* in young (3-week-old) and aging (13-week-old) plants (Figure 2A & B, respectively). The highest expression levels of these three genes were found in flowers and siliques of 13-week-old plants, in which the abundance of *pri-miR840* in the reproductive organs was about ten-fold higher than in the vegetative organs (Figure 2B). The expression levels of *pri-miR840* in rosettes of 3-week-old seedlings and 13-week-old plants were comparable, whereas *PPR* and *WHY3* showed a decreased (~ −5-folds) and enhanced (~ +2.5-folds) expression with increasing age, respectively (Figure 2A & B).

**Figure 2.**
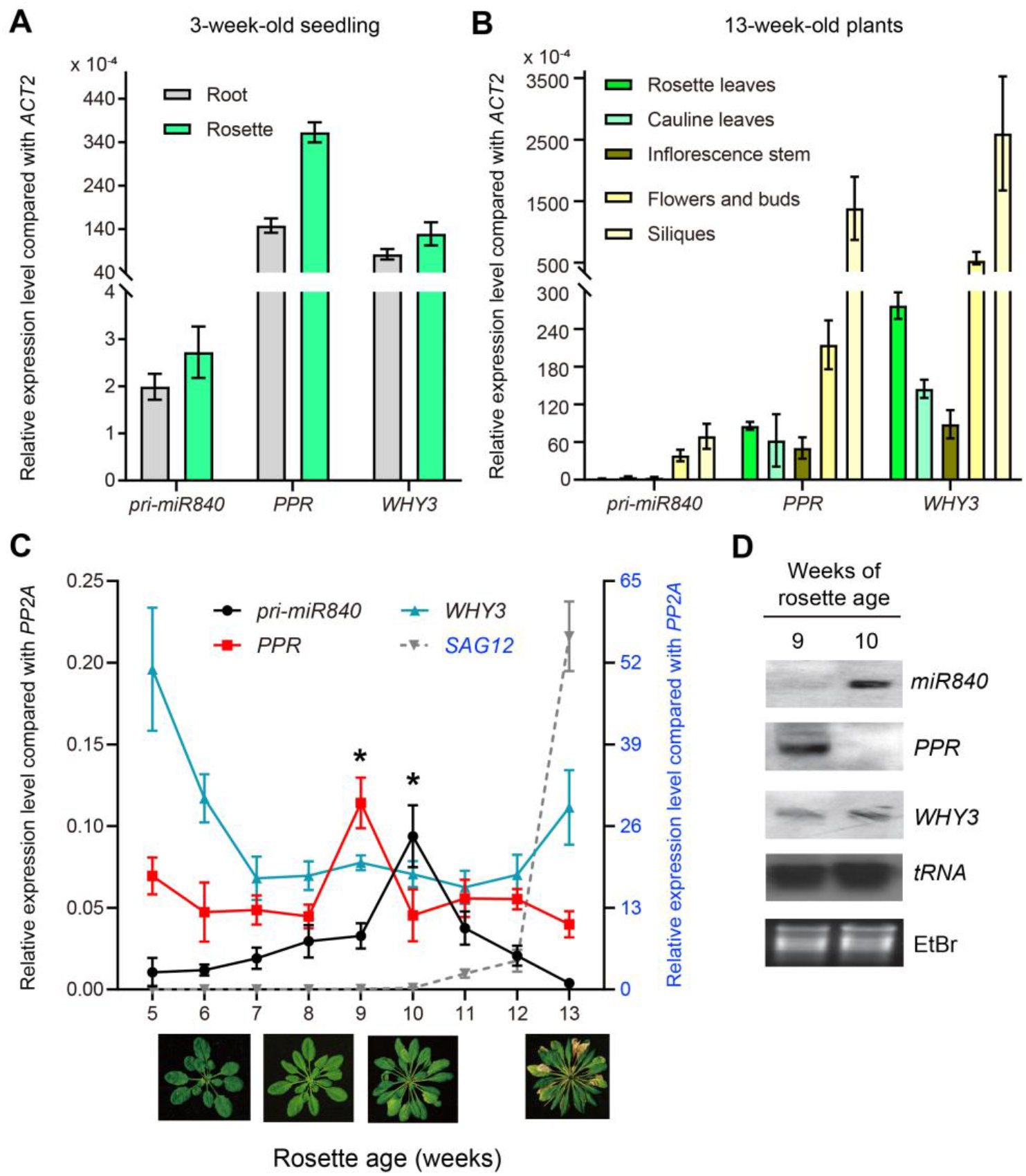
Expression of *MIR840, PPR* and *WHY3* during development and senescent process. **(A)-(B)** Tissue-specific expression in rosette and root of 3-week-old seedlings and in 13-week-old plants, respectively. **(C)** Age-dependence expression profile in rosettes. The senescence-associated gene *SAG12* was used as a molecular marker of senescent stage of Arabidopsis plants. **(D)** Northern blot of miR840, *PPR* and *WHY3* transcripts in rosettes of 9- and 10-weeks when senescence is set. EtBr: ethidium bromide staining.

To monitor the transcript profiles of miR840 and *PPR* and *WHY3* during plant development and aging, we weekly sampled rosettes from week 5 to week 13. During this period, week 10 marked the senescent initiation stage under our experimental conditions, with activation of a senescence-associated marker gene *SAG12* (Figure 2C). Both *PPR* and *WHY3* displayed an antagonistic expression pattern from week 9 on. Interestingly, this time point marked the highest expression of *PPR* throughout all developmental stages, preceding a similar expression profile of miR840 being one week delayed and coincident with the senescent initiation (Figure 2C). The transcript profiles of mature miR840 as well as of *PPR* and *WHY3* in 9- and 10-week-old rosettes were additionally confirmed by Northern blot hybridization (Figure 2D). Furthermore, we also determined the levels of mature miR840 as well as *PPR, WHY3* and *SAG12* transcripts during the aging of rosette leaves (with leaves from different positions) and in 4-sectioned leaf segments from yellowish tip to green base of the single 7^th^ leaf of 11-week-old plants (Figure S1 and S2, respectively). Similarly, a significant elevation in miR840 abundance was associated with the onset of leaf senescence as indicated by an up-regulation of *SAG12* (Figure S1C). While *PPR* expression could be negatively correlated with miR840 expression during plant aging, this was not true for *WHY3* (Figure S1B and S2C). These data suggest a possible involvement of miR840 in plant senescence regulation.

### Loss-of-function and gain-of-function analysis indicates a crucial role of miR840 in plant senescence regulation

Two homozygote T-DNA insertion lines, *SALK_038777* and *SAIL_232_F08* were employed for further analysis. Both lines can be considered as *MIR840* mutants inserted at promoter position (Figure 3A and Figure S3), but also disrupt the *PPR* ORF. The T-DNA insertion at position −767 bp (*SALK_038777*) reduced the miR840 level about 95 folds as compared with the WT, whereas the insertion at −384 bp *(SAIL 232_F08)* drastically enhanced miR840 expression up to approximately 45-fold as revealed by northern blot and qRT-PCR analysis from rosettes harvested at the onset of plant senescent stage (week 10). Thus, we considered *SALK_038777* as a miR840-knockdown and *SAIL 232_F08* as an overexpression line for this study (Figure 3A). However, it is not yet clear how miR840 expression is affected by the T-DNA-insertions (Figure S3).

**Figure 3.**
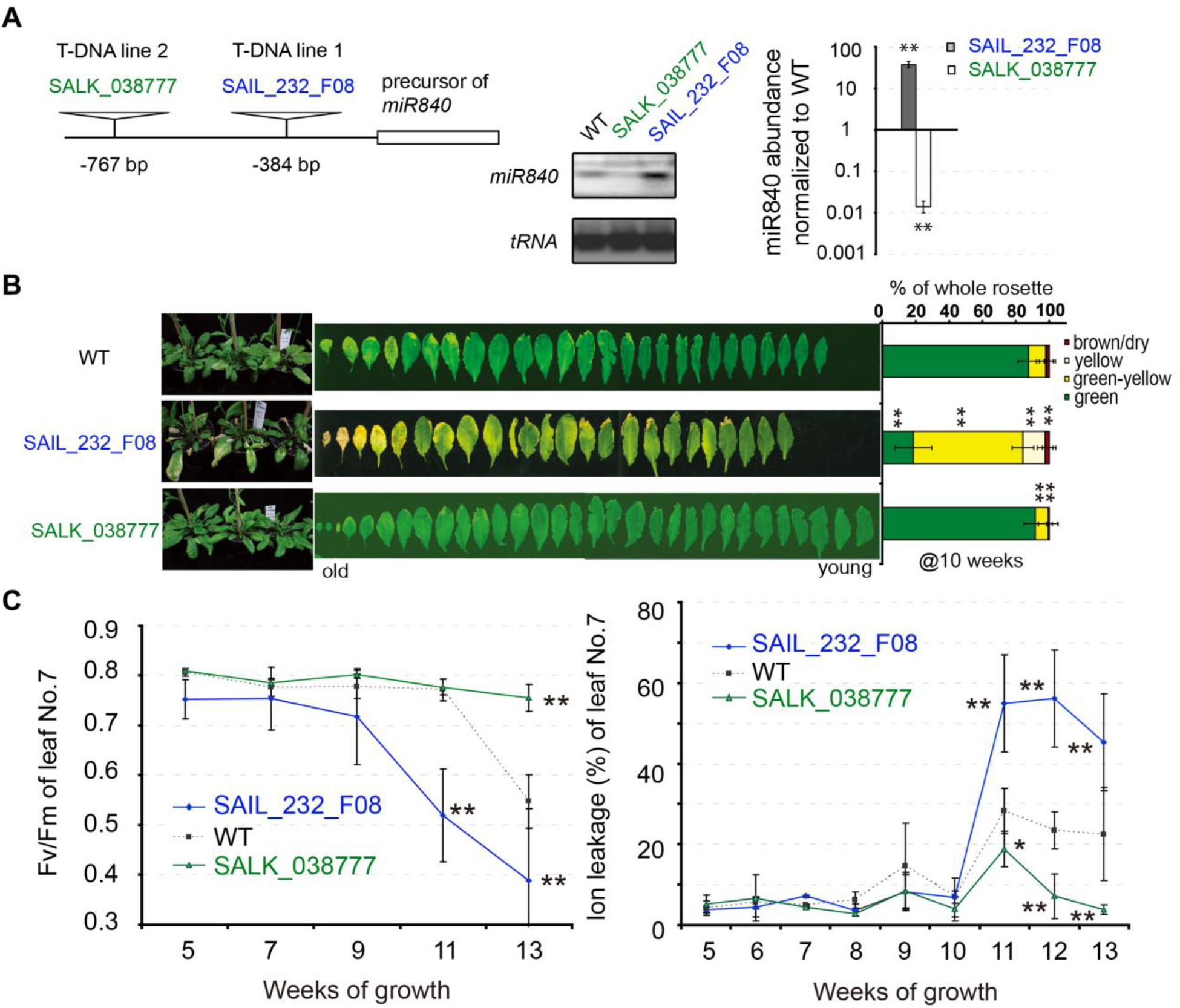
Phenotypical characterization of the two T-DNA mutant lines of Arabidopsis. **(A)** Positions of T-DNA insertion and the effect on miR840 expression of the two *mir840* mutations. **(B)** Senescent phenotypes comparing with WT plants. Twelve 10-week-old plants were measured for leaf color scoring under 8 h illumination condition (mean ± SD). Representative photos are shown. **(C)** Relative photochemical efficiency of photosystem II (Fv/Fm) (left panel) and ratio of the membrane ion leakage (right panel) in rosette leaf No.7 were determined at different developmental and senescent stages. The data are mean ± SD of twelve (for Fv/Fm) or five (for ion leakage) independent measurements. * *P* < 0.05, ** *P* < 0.01.

Phenotypically, both mutants displayed contrasting leaf development and senescence onset. The *SALK_038777* with lower miR840 expression showed a stay-green phenotype even superior to the WT, whilst the *SAIL_232_F08* with miR840 overexpression exhibited a strong early senescence-like phenotype (Figure 3B). Qualitative and quantitative determination of senescence related parameters, such as leaf yellowing (Figure 3B), photochemical efficiency of photosystem II F_v_/F_m_ and leaf ion leakage (Figure 3C) suggested that miR840 has a strong impact on plant development at the later stages (at about 10 weeks). The observed phenotypes were stable and could be confirmed by further measurement with up to 7^th^ generation of the mutant plants (Table S1).

To verify the function of miR840 in plant development and senescence observed from the T-DNA insertion lines, we further generated transformants in Arabidopsis plants ectopically expressing either the *pri-miR840* and a tandem antisense (target) mimicry of miR840 *(9x miR840am)* (Figure 4A; Figure S4A and S4B). By stem-loop RT-qPCR we found the *pri-miR840* OE lines accumulating 289-to 1100-fold more miR840 transcripts when compared with WT, while in the antisense mimicry lines mature miR840 decreased 3-10 fold of WT levels (Figure 4B). Phenotypic analysis revealed that the overexpression of miR840 was indeed associated with early senescence, whilst the knockdown of miR840 by antisense mimicry delayed plant senescence, as indicated by measurements of leaf yellowing, Fv/Fm index, total chlorophylls and total carotenoids contents (Figure 4C and 4D).

**Figure 4.**
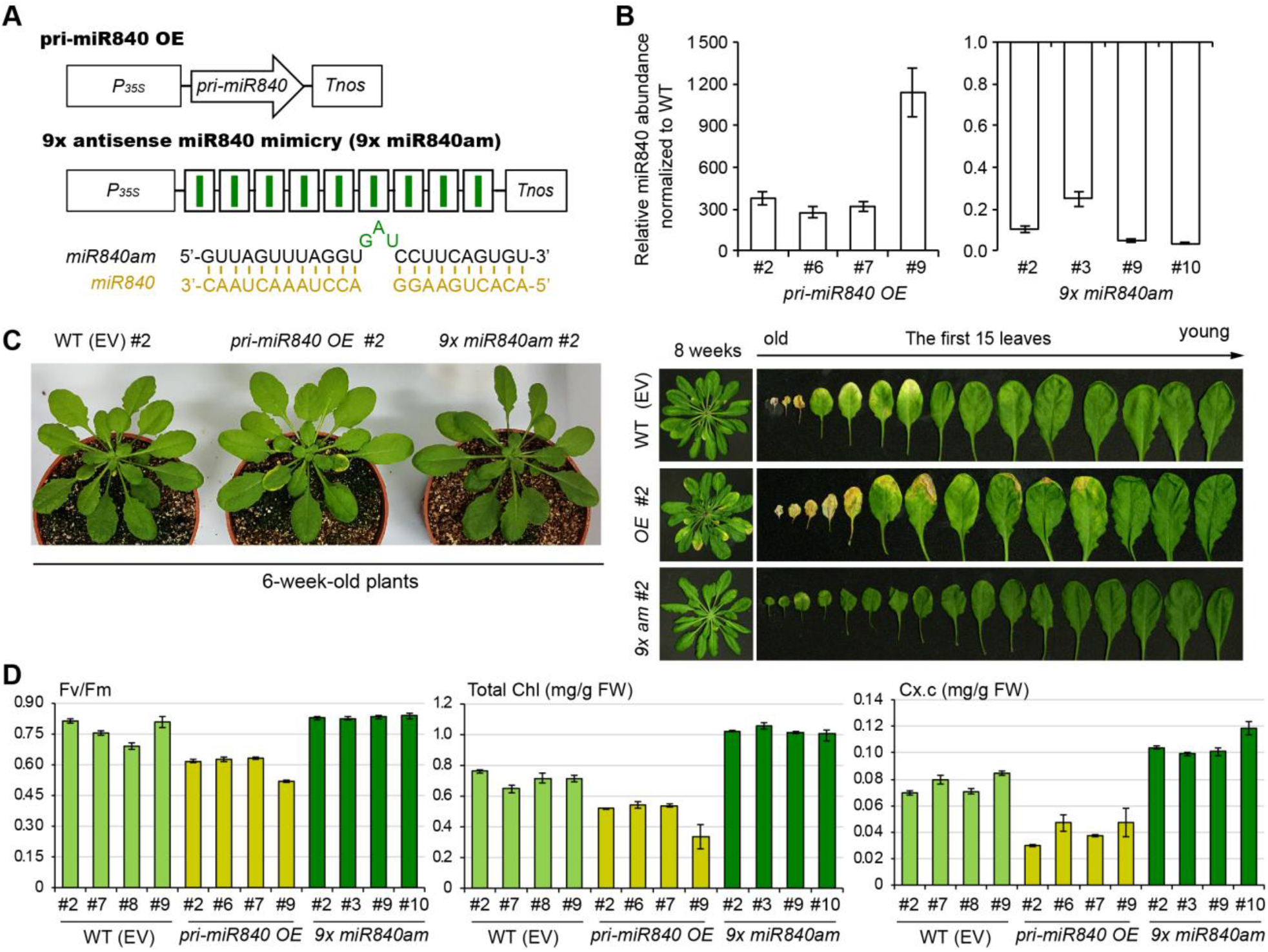
Senescence-related phenotyping of *pri-miR840 OE* and *9x miR840am* transgenic plants. **(A)** Schema of the expression cassettes for transgenes *pri-miR840 OE* and *9x miR840am. P35S, CaMV35S* promoter; *Tnos, NOS* terminator. **(B)** Determination of miR840 accumulation in rosettes of independent transgenic lines, showing fold-change over WT plants. **(C)** Senescent phenotypes of 6- and 8-week-old WT, *pri-miR840 OE* and *9x miR840am* transgenic plants. Representative 6-week-old plants showing early senescence in the OE transgenics as compared to the others (left panel). Representative 8-week-old plants with ordered rosette leaves were displayed (right panel). **(D)** Measurement of Fv/Fm value (left panel), total chlorophyll content (middle panel) and total carotenoids content (Cx.c) (right panel) in the 4^th^ rosette leaf of 6-week-old plants. Data represented mean ± SD (n =5).

The phenotypical differences observed in the T-DNA KD and OE mutants of miR840 were correlated with senescence-associated gene expression as demonstrated by RT-qPCR of 50 senescence-related genes (Figure S5) known to be directly involved in plant senescence or cell death and DNA damage/repair processes. The results showed that the expressions of these genes were differentially affected in the miR840 mutants as compared to WT, albeit to different degrees (Figure S5B). In the miR840-overexpression mutant *SAIL_232 F08,* a strong increase in gene expression was observed for *WRKY53, WRKY33, SIRK, SAG101, SAG12,* and *PDFs.* Consistently, a significant decrease of the gene expression was detected for *WRKY53*, *WRKY33*, *SAG101* and *PDFs* genes in the miR840-knockdown mutant. Interestingly, the expression of *PR1* gene was increased in *SALK_038777* but decreased in *SAIL_232_F08* plants, hinting at a possible crosstalk between senescence and the plant defence response. Moreover, the expressions of *SPO11-2* and *RAD52,* two genes functioning in double-stranded break (DSB) related DNA damage and repair mechanism, were up-regulated in *SAIL_232_F08* OE but down-regulated in *SALK_038777* KD mutants (Figure S5B). Taken together, we conclude that miR840 represents a master-regulatory microRNA in reprogramming cellular pathways to enhance or initiate plant senescence.

### MiR84O induces atypical cleavage of its target mRNAs

miR840 is transcribed in the same orientation as *PPR* and both 3’-UTRs of *PPR* and *WHY3* are predicted targets of miR840* and miR840, respectively (Figure 1A). To detect the miR840-target sites in *PPR* and *WHY3* transcripts, we performed a 5’-RLM-RACE experiment using total RNA extracted from rosettes of WT plants at the senescence onset stage when miR840 was highly accumulated. The efficiency of reverse transcription reaction of the adaptor-ligated cDNA was exemplified by amplification of a reference gene *AtUBQ13* and an extremely lowly expressed gene *AtSUC7* (Figure S6A), whereas miR164-guided cleavage of *NAC1* transcript (Guo et al., 2005) served as a positive control (Figure S6B). Surprisingly, sequencing of the cleaving products of both *PPR* and *WHY3* transcripts revealed their target sites outside the conventional admitted region of the miR840 complementary sequences. In *WHY3* it was found to be 9 bases downstream (close to the polyA-tail) of the miR840 binding sequence, whereas in *PPR* transcript it was located 22 bases downstream of the miR840* pairing region (Figure S6B and S7). Such noncanonical cleavage events have also been reported in other G3A type microRNAs, such as miR844-targeted *CDS3* (Lee et al., 2015).

The target sites were further verified using an *in vitro* mutagenesis experiment. A reporter plasmid was constructed by cloning synthetic oligonucleotides, resembling the 78-nt-region carrying the miR840-targeting sequence or bearing a mutated targeting site, into a *CaMV35S*-promoter-driven GUS-expression vector (Figure 5A). A 35S promoter-driven *pri-miR840* expression plasmid was used as an effector and transferred, in combination with the respective reporter plasmid, into *Nicotiana benthamiana* by Agrobacterium-mediated infiltration (Figure 5B). Single infiltration of the reporter plasmid showed slightly decreased *GUS* transcription when it was fused with the original *WHY3* or *PPR* 3’-UTR as compared to that without fusion R (0). But both fusion constructs did grant strong GUS staining (Figure 5C). After mutation of the target sites of miR840, the *GUS* transcripts were further reduced in both fusion cases and their GUS staining signals became moderate as compared with R (0) (Figure 5C). Nevertheless, co-infiltration with the effector construct caused a drastic reduction in *GUS* transcript levels as well as the GUS staining signals to ~ 20% of the respective single construct with the original *WHY3* or *PPR* 3’-UTR. However, such reduction effects caused by co-infiltration with miR840 did not ocurr if the *PPR* or *WHY3* target site were mutated, which confirms that the determined target sites are no artefacts (Figure 5C). Following these data, we conclude that miR840/miR840* targets the overlapping 3’UTR region of *WHY3* and *PPR* transcripts, respectively, but with atypical target sites outside the respective pairing region (Figure S7).

**Figure 5.**
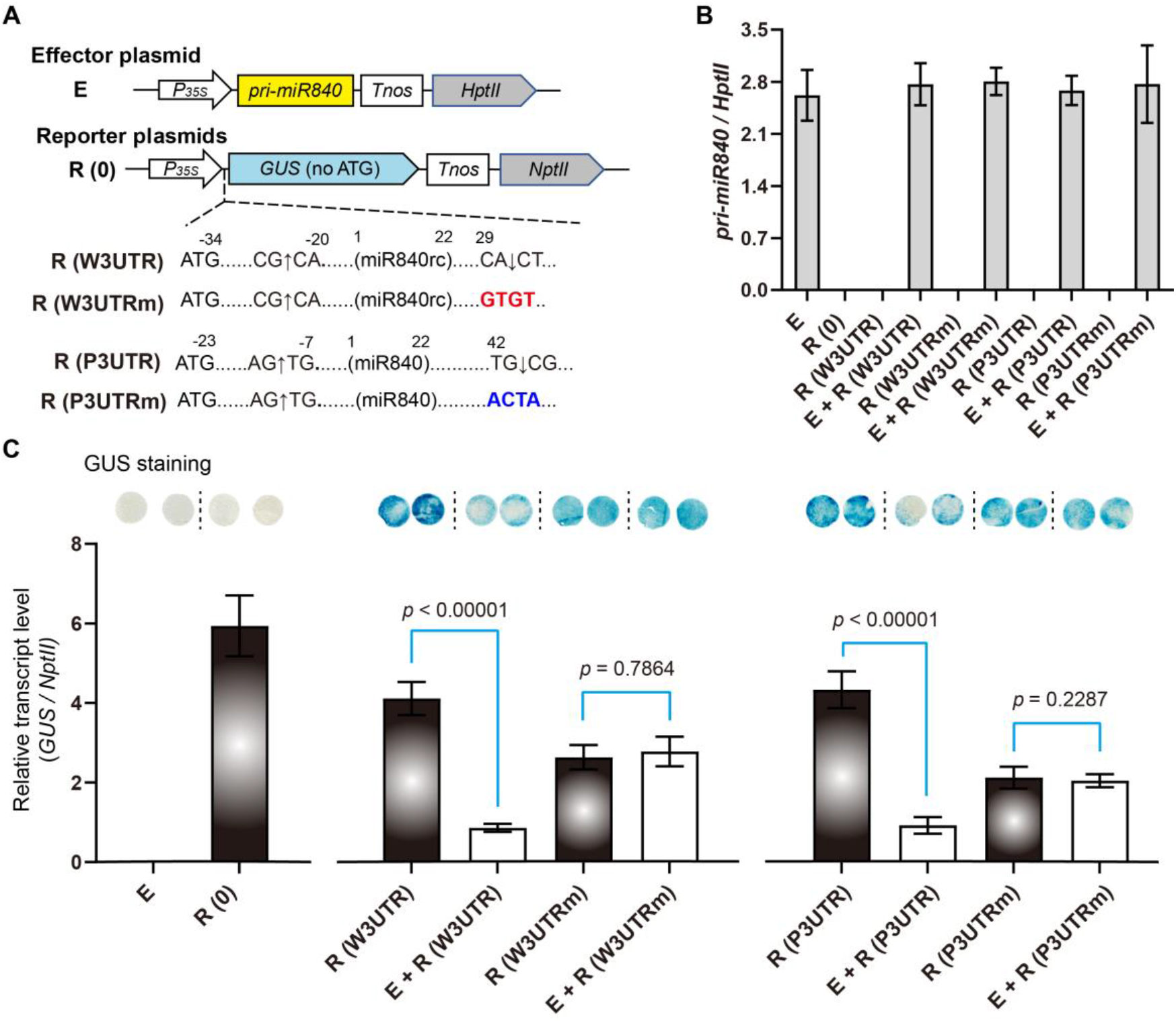
A GUS report assay demonstrating a site-specific dependency in miR840-guided transcript cleavage. **(A)** Effector and reporter plasmids designed in the *N. benthamiana* transient assay. The ATG-less *GUS* cDNA is transcribed downstream of and in frame with a 78 bp short sequence containing the specific *miR840* cleavage sites in the *3’UTR* region of *PPR* and *WHY3* transcripts, respectively. Nucleotide substitutions are highlighted in red and blue, respectfully. **(B)** qRT-PCR detection of *pri-miR840* abundance at three days post infiltration, using the plasmid-encoded *NptII* as an internal reference. **(C)** Transcript level and GUS staining at three days post infiltration. Statistical significance was determined using a paired Student’s t-test with alpha = 0.05; the adjusted p-value is shown above the data (n = 5).

### MiR840 targets *PPR* post-transcriptionally but *WHY3* is inhibited translationally

In the next step, we further checked how miR840/miR840* affects *PPR* and *WHY3* expression *in planta*. Unexpectedly, the *WHY3* transcript level was not affected in the two T-DNA lines, *SALK_038777* and *SAIL_232_F08* as determined by RT-qPCR analysis (Figure 6A). In contrast, the *PPR* expression was conversely regulated as expected in both the two T-DNA mutant lines (Figure 6A) as well as in the miR840 mutants *pri-miR840 OE* and *9x miR840am* (Figure 6B). These results indicated a strong negative correlation between the *PPR* expression and the miR840 levels.

**Figure 6.**
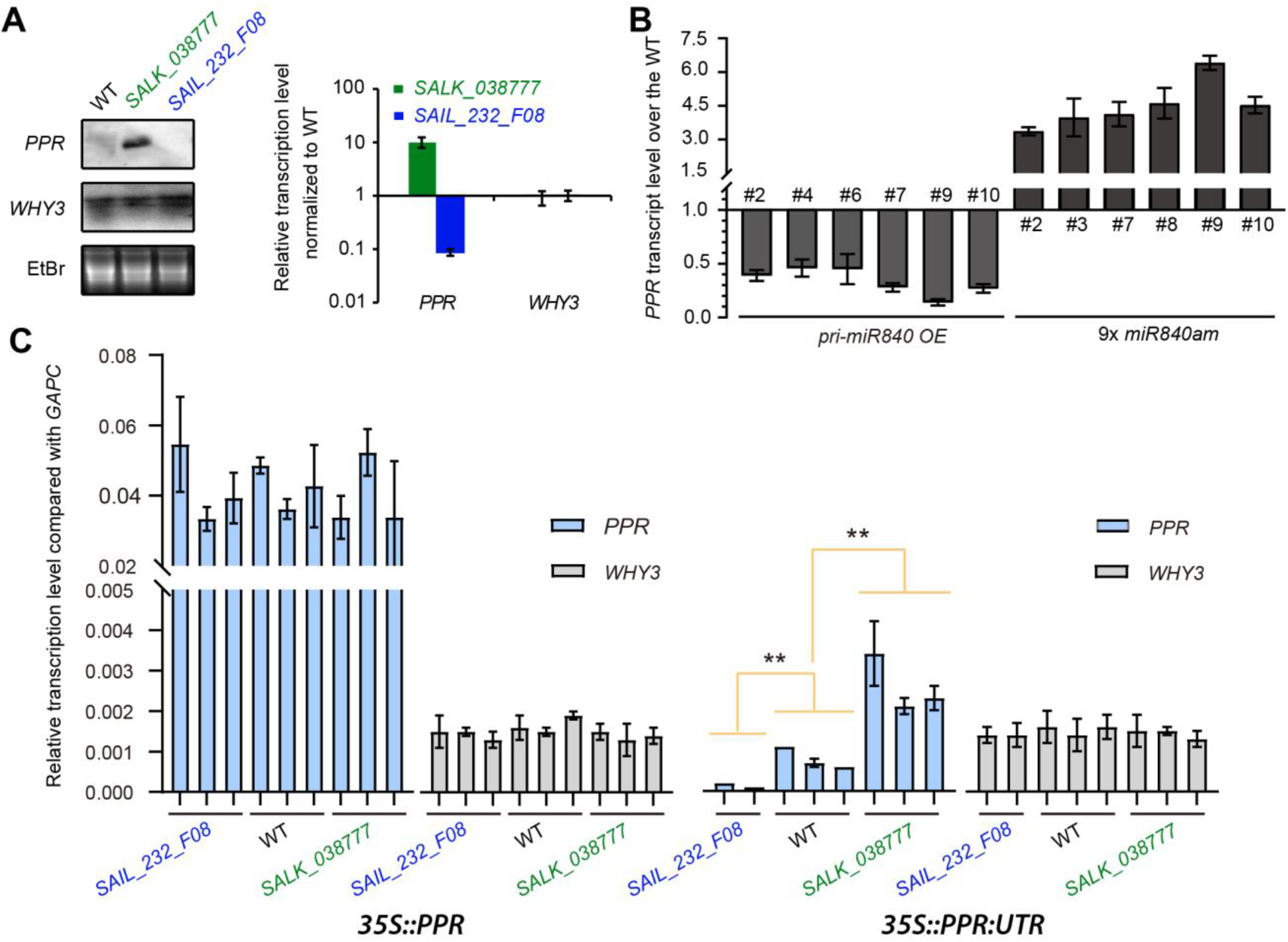
*PPR* expression is targeted post-transcriptionally by MiR840. **(A)** Northern blot and qRT-PCR analysis of *WHY3* and *PPR* transcripts in miR840 gain-of-function mutant *(SAIL_232_F08)* and loss-of-function mutant *(SALK_038777)* and in WT plants. **(B)** Fold-change in transcript abundance of *PPR* in transgenic lines of *pri-miR840 OE* and *9x miR840am*, comparing to the WT. **(C)** mRNA level of *PPR* and *WHY3* in rosettes of transgenic lines over-expressing *PPR* CDS either with or without its 490-bp 3’UTR in WT or miR840 mutant background. Data are mean ± SD (n =3). ** *p* < 0.01.

To ascertain that PPR is indeed post-transcriptionally regulated by miR840*, we overexpressed *PPR* with or without the 3’UTR region in the two T-DNA lines, as well as in WT. Again, the *WHY3* expression level was not affected in all transgenic plants (Figure 6C) and a strong reduction of *PPR* transcript levels was found in transformants harbouring the 3’UTR-containing construct, while the reduction of *PPR* transcripts was protected in transformants harbouring the 3’UTR-less construct (Figure 6C). Moreover, in transgenic plants expressing the 3’UTR-containing construct, the *PPR* expression levels varied depending on the genetic backgrounds, generally negatively correlating with miR840 expression (Figure 6C).

Even though *PPR* and *WHY3* are both targets of miR840, the *WHY3* transcript level was not affected in the two T-DNA lines, *SALK_038777* and *SAIL_232_F08* (Fig. 6A), We assumed that the miR840 might then interfere with the WHY3 translation. To test this hypothesis, we performed western blot analysis using an antibody against a specific WHY3 peptide (Fig. 7A). As a result, the protein level of WHY3 in the miR840-OE (*SAIL_232_F08*) was much lower than that in both the WT and the miR840-KD *(SALK_038777)* (Figure 7A). To verify the miR840 pairing effect, we deployed three constructs in either the WT or the two T-DNA mutant backgrounds to express estradiol inducible WHY3-luciferase fusions with the WHY 3’-UTR sequences *(UTR)* or its cleavage-site-mutated form *(UTRm)* in the pMDC7 vector (Figure 7B). The WHY3-luciferase without WHY 3’-UTR (CK) served as a control. Induced expression of the recombinant fusion proteins was monitored during 1 – 48 h post induction (hpi) in the WT plants transformed using the *UTR* construct (Figure 7C, left panel). Without the induction, the fusion protein expression levels were very low in the *UTR*-harboring plants regardless of the genetic backgrounds, whilst at 24 hpi the WT plants and the miR840-KD showed strong signals in the western blot (Figure 7C, right panel), but not in miR840-OE mutant lines, suggesting an translationally inhibitory effect of miR840.

**Figure 7.**
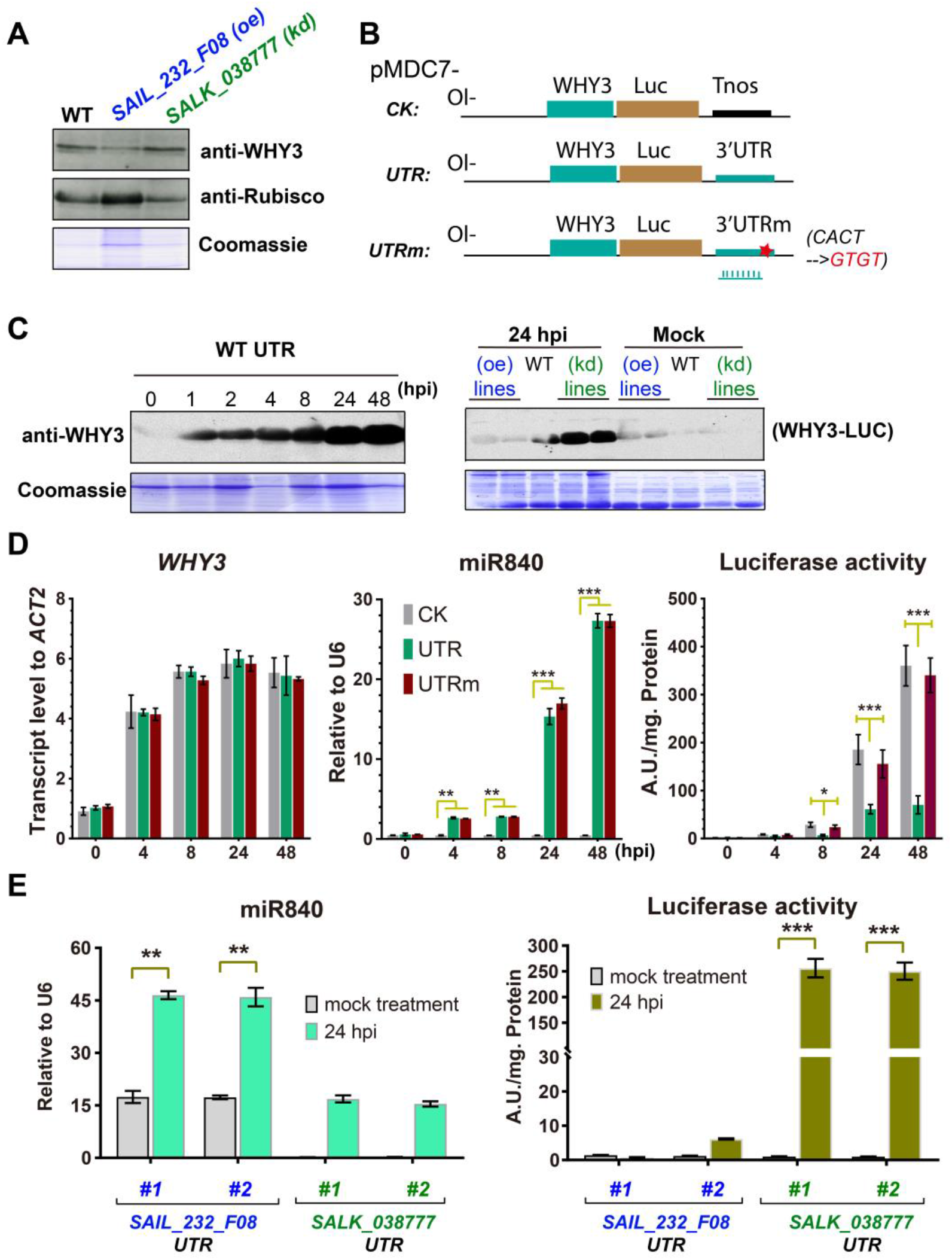
*WHY3* expression is translationally inhibited by MiR840. **(A)** Western blot showing WHY3 protein level in *SAIL_232_F08, SALK_038777* and WT plants. The anti-Rubisco and Coomassie staining were used for loading control. **(B)** A schema showing the estradiol induced constructs for *WHY3-LUC* in frame fusions based on the pMDC7 vector. *CK*: no additional sequence was added after the fusion; *UTR:* the 560 bp WHY3 3’UTR sequence was inserted downstream of the fusion; *m-UTR:* the 560 bp WHY3 3’UTR sequence with mutated miR840 cleavage site (nucleotides in red). **(C)** Time-course of fusion protein accumulation after estradiol treatment in stably transformed WT plants overexpressing the *UTR* construct (left panel), and the 24-h-induced fusion protein levels in the three transgenic plants stably expressing the *UTR* construct (right panel). Coomassie staining is used as a loading control. Note that the fusion protein is repressed in the gain-of-function miR840 mutant **(D)** Quantification of miR840 and *WHY3* transcript level, as well as luciferase activity in transgenic plants of the WT background after treatment with estradiol. Note that the luciferase activity is blocked in plants with *UTR* construct but not in the *UTRm* and *CK* plants. **(E)** Correlation of miR840 abundance (left panel) and luciferase activity (right panel) using *UTR* transgenic plants of the two miR840 mutants. Data of two independent lines from each are shown. Data are given as mean ± SD of three biological replicates. * *p* < 0.05; \*\**p* < 0.01; *** *p* < 0.001. A.U., arbitrary unit.

*WHY3* transcript levels in WT plants transformed with the three inducible constructs were induced up to 2 – 6 folds by estradiol treatment, and comparable among the two constructs, *UTR* and *UTRm* (Figure 7D). However, a significant increase in miR840 levels after estradiol treatment was also detected in both *UTR-* and *UTRm-*expressing plants, but not in the *CK* plants (Figure 7D). In both cases, it was probably caused by a homolog seed sequence of 3’-end miR840 located at the *WHY3* 3’-UTR (4 bp difference out of 22 bases to miR840, Figure 1B) that could be mis-amplified by the stem-loop qRT-PCR. Therefore, it could be considered as a background noise signal. Nevertheless, the activity of firefly luciferase was gradually increased in all the three types of *UTR* plants after estradiol induction, but much more pronounced in plants harboring *Tnos* and *UTRm* constructs than in those with *UTR* (Figure 7D). Thus, under WT backgrund, expressed fusion with the original 3’-UTR of WHY3 resulted in inhibition of protein synthesis, whilst this inhibitory effect was dismissed, when the miR840-targeting site at the 3’ -UTR was mutated.

We further compared the 3’UTR effect on ectopic expression of firefly luciferase (LUC) activities in the two T-DNA insertion lines with endogenous accumulated or depleted miR840 level. Again, detection of miR840 via stem-loop qRT-PCR posed a background signal in both lines, quantitatively similar to that in the *UTR*- or *UTRm*-expressing WT shown in Figure 7D. Yet, the *SAIL_232_F08* plants did have noticeably higher miR840 accumulation than the other line (Figure 7E, left panel). As expected, in the miR840-*OE SAIL_232_F08* LUC activities were lowly detectable regardless of induced conditions, indicating that the endogenous miR840 could have inhibited the induced expression of the WHY3-LUC fusion (Figure 7E, right panel). On the contrary, in *SALK_038777* after estradiol treatment, LUC activity was markedly increased (Figure 7E). Theses results supported for an inhibitory effect of the fusion reporter by miR840.

Taken together, we demonstrate that both *WHY3* and *PPR* are targeted by miR840/miR840* at their 3’-UTRs, respectively. Whereas pairing of miR840* to the *PPR* mRNA leads to its degradation, miR840 inhibits the protein synthesis of WHY3.

### Both *PPR* and *WHY3* are regulators of senescence with common downstream genes

To investigate the mode of action of miR840-*WHY3* and miR840-*PPR* in plant senescence, a T-DNA insertion line of *WHY3* (*Salk_005345C*) was obtained, in which the T-DNA insertion at −140 bp upstream of its start code (ATG) (Figure S4C) resulted in a 20-fold decrease in *WHY3* expression (Figure 8A, left panel insert; Figure S8B and C). This mutant line was assigned *kdwhy3*. In parallel, transgenic plants using a *PPR* antisense construct (designated as *appr*) was generated, in which *PPR* expression was strongly depressed as compared to WT (Figure S8A and C). However, neither *kdwhy3* or *appr* plants showed accelerated senescent phenotypes as compared to WT (Figure S8D to G), suggesting they worked in concert to initiate the onset of senescence. Therefore, we further generated double mutant by crossing of *kdwhy3* x *appr.* The homozygote double mutant plants *kdwhy3 appr* selected from a F3 generation were employed for further analysis. As assumed, the double mutant showed now an early senescence phenotype similar as that observed in the miR840-overexpression lines (Figure 8A and B; Figure 3B and C).

**Figure 8.**
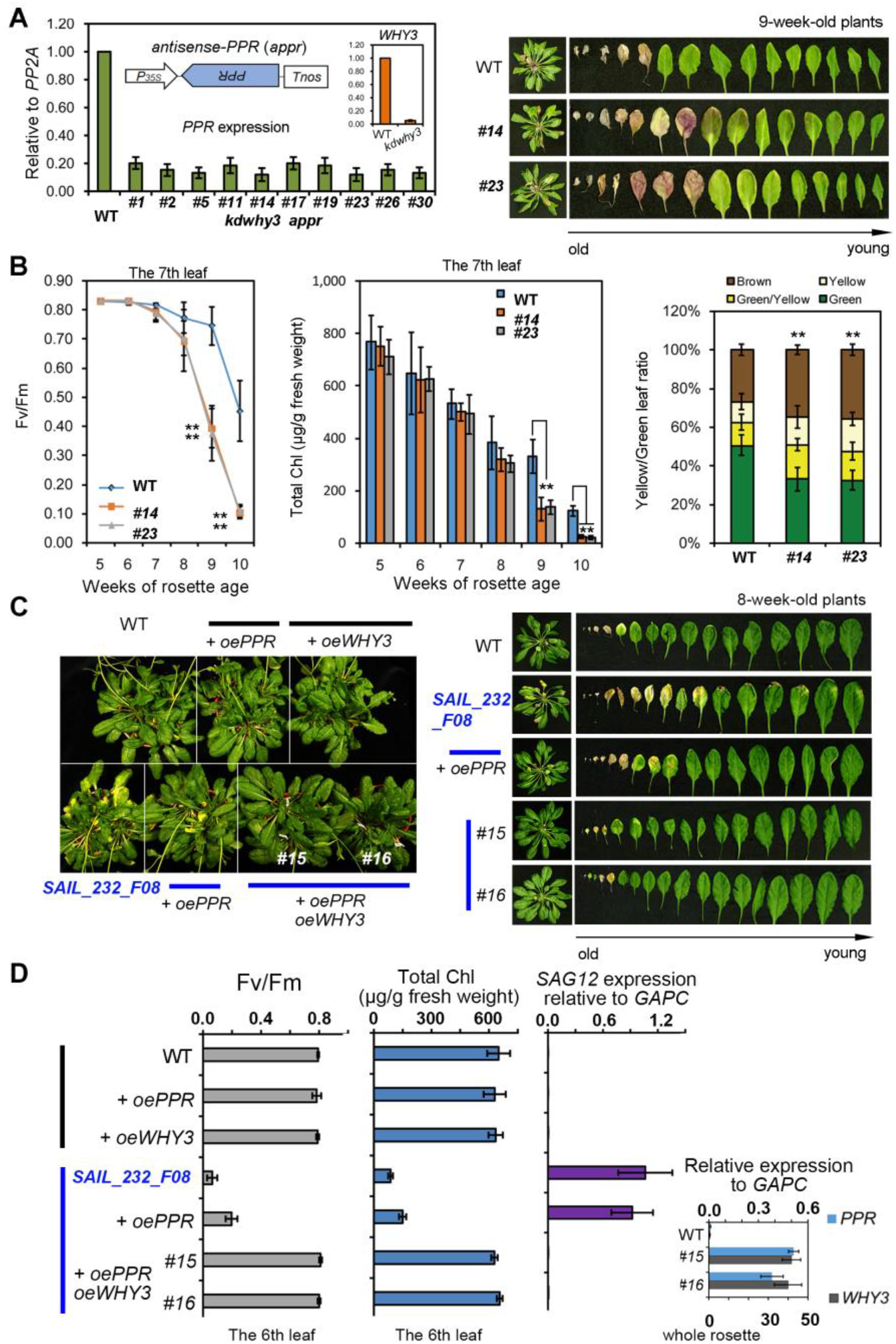
Double-knockdown mutants of *PPR* and *WHY3* show similar early-senescence phenotype with the gain-of-function miR840 mutant to a less extent. **(A)** A *kdwhy3 appr* double mutant was generated by stable transformation of the *kdwhy3* mutant (T-DNA insertion line, *SALK_005345C,* Figure S4C) with a *PPR* antisense construct *(appr).* Representative photos of the rosette leaves are shown (right panel). **(B)** Determination of relative photochemical efficiency of photosystem II (Fv/Fm) (left panel) and total chlorophyll content in rosette leaf No. 7 (middle panel), as well as of proportion of leaf coloring of the rosette leaves (right panel). The value represents mean ± SD of 12 independent measurements for Fv/Fm and Chl content, and 15 for yellow/green leaf ratio. Significant difference: * *p* < 0.05, \*\**p* < 0.01. **(C)** Phenotype analysis of stable transgenic plants overexpressing *PPR* (*oePPR*) or *WHY3* (*oeWHY3*) in the WT background and *oePPR* alone or together with *oeWHY3* in the miR840 gain-of-function mutant *SAIL_232_F08*. Note that the early senescence phenotype of the *SAIL_232_F08* can be restored to that of WT by co-expression of *PPR* and *WHY3.* **(D)** Quantification of relative photochemical efficiency of photosystem II (Fv/Fm) (left panel), and total chlorophyll content (middle panel) in the 6^th^ rosette leaf, as well as *SAG12* gene expression in the same plants as in (C). The insert shows transcript levels of *PPR* and *WHY3* in WT plants and the transgenic plants.

Moreover, to rescue the early senescence phenotype in *SAIL_232_F08* OE mutant, we stably overexpressed *PPR* alone (*oePPR*) or together with *WHY3* (*oePPR oeWHY3*) in this background (Figure 8C and D). Only transgenic plants overexpressing both *PPR* and *WHY3* genes could rescue the early senescence of the mutant comparable to the WT plants, whilst overexpression of the *PPR* alone had only a very weak rescue effect (Figure 8C and D).

The double mutant *kdwhy 3appr* does not only resemble the early senescece phenotype of the miR840-overexpression plants, but also displayed a similar expression profile of downstream genes known to be involved in senescence, cell death and DNA damage repair (Figure 9A to C, and Figure S5). Several of these genes also showed similar expression patterns in the single *kdwhy3* or *appr* plants (Figure S9D and E). Notably, similar regulation of *CAMTA3, AHK3* and genes coding for a pyruvate decarboxylase (*AT5G01320*) and a glutamine synthase (*GLN1*) were found among *kdwhy3*, *appr* and *kdwhy3 appr* transgenic plants as well as in the miR840-overexpression lines (Figure S9D, E and Figure S5). Besides that, *arginine-tRNA protein transferase gene (DLS1), RLP27, ANAC053, COR15B, NEET, AT4G22620, WRKY70* and *APG7* genes shared the similar expression patterns in *appr* and the the miR840-overexpression lines (Figure S9E; Figure S5), and *copper amine oxidase (AT4G12290), COR78, ARF2, UBA2A, ANAC092*, *ARF1*, *ANAC029*, *COR15A*, *WRKY70*, *MYB34*, *LEA hydroxyproline-rich glycoprotein* (*AT1G17620*), *B12DP* and *Copia-like retrotransposon* (*AT5G35935*) genes exihibted similar expression patterns in *kdwhy3* and the miR840 overexpression lines (Figure S9E; Figure S5).

**Figure 9.**
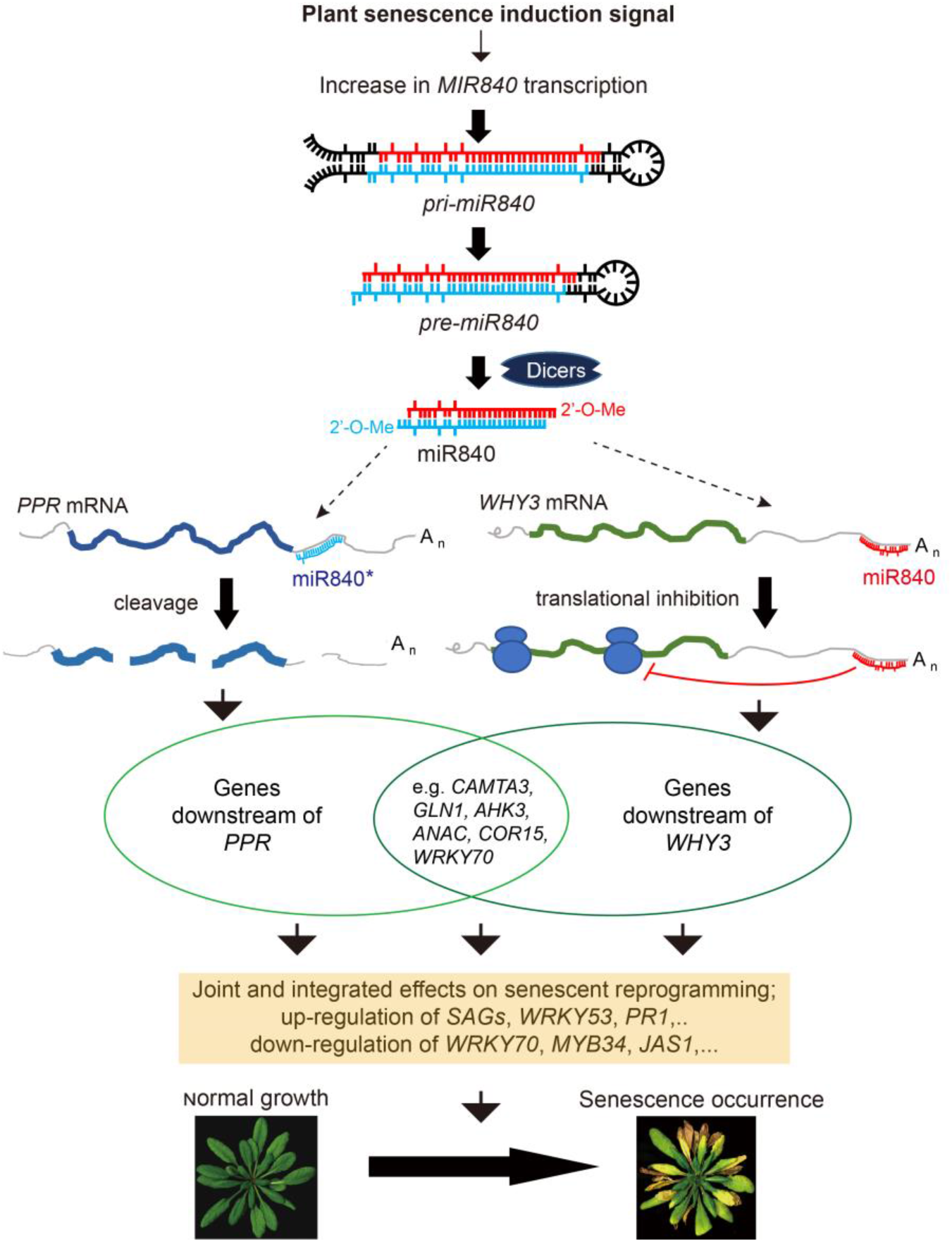
A working model showing how miR840 regulates plant senescence in Arabidopsis. Increased expression of *MIR840* may be activated by developmental or environmental signals. Processing of *pri-miR840* and generation of mature miR840 involve many factors but are controlled by *DCL1, DCL4* and to a less extent *DCL2.* On the contrary, *DCL3* is not required for miR840 production. Accumulated miR840 or its star strand is then loaded on ARGONAUTE proteins as part of the RISC (RNA-induced silencing complex) and target to mRNAs of *WHY3* and *PPR* by pairing with the complementary region of the respective transcripts. Both miR840 and miR840* are able to bind to *WHY3* transcripts with perfect match, and to *PPR* transcripts with four mis-matches, and the binding leads to the cleavage at specific sites downstream of the pairing regions, which are considered as nonconventional. Whereas *PPR* transcripts are reduced by miR840-guided degradation, WHY3 protein synthesis is inhibited by a miR840-mediated mechanism. Target repression by miR840 may include other unproven genes, however, concurrent knockdown of *WHY3* and *PPR* is sufficient to mimic the early senescence phenotype of gain-of-function miR840 mutations. Down-regulation of *WHY3* and *PPR* provokes up-regulated expression of a set of senescence-associated genes (*SAGs*), which in turn triggers the initiation of senescent progression. The miR840-*PPR*/*WHY3* regulatory pathway of plant senescence seems to be a new module limited to plants of the genus Arabidopsis. arrow with slim line: effect or regulation, arrow with fat line: transformed process, arrow with broken line: multiple steps, line with stop: inhibition

We conclude that both *PPR* and *WHY3* are targets of miR840-mediated senescence pathway, and simultaneous repression of both genes by miR840 reprograms the expression of a subset senescence-related genes, consequently leading to the onset of plant senescence (Figure 9).

## Discussion

### miR840 represents an evolutionary young microRNA with a special targeting configuration

The special genomic arrangement of the *miR840* locus at the convergent region between 3’UTRs of *PPR* and *WHY3* makes it possible that miR840 may be expressed from its own promoter or generated from 3’UTR of a *PPR* transcript. Our results indicate that miR840 is mainly produced by Dicer-dependent pathways. Among the four members of the Dicer family in Arabidopsis, DCL1, 4 and 2 are contributing to the processing of the mature miR840 (Figure 1). Although DCL1 can be considered as the major dicing enzyme for miRNA biogenesis (Reinhart et al., 2002), DCL4 is also involved on generation of evolutionary young miRNAs (Rajagopalan et al., 2006). Both DCL4 and DCL2 preferentially produce 21- and 22-nt siRNA from endogenous or viral or transgene, respectively (Borges and Martienssen, 2015; Taochy et al., 2017). While the effect may not be significant, it is interesting here that miR840 (22-nt) production is also partially dependent on DCL2, but a detailed analysis on the RNA species produced by different DCLs is necessary in helping to unveil the underlying mechanism.

Both strands of the duplex miR840 can bind to the mRNAs of *WHY3* or *PPR*, yet the former pairings yield perfect matches but the latter ones consist of 4 mismatches (Figure S7). Such configuration also means that targeting to *WHY3* and *PPR* by mature miR840 may involve an unconventional mechanism, because perfectly matching between miRNA and target sequence is usually associated with cleavage. However, we observed translational repression as the predominant mechanism to prevent WHY3 expression, and cleavage for the PPR transcript despite of the 4 mismatches. Our data revealed furthermore that miR840-guided target sites in both *WHY3* and *PPR* transcripts located outside of the predicted pairing regions (Figure S6), with two distinct consequences – translational inhibition for *WHY3* and mRNA degradation for *PPR*. Similar cleavage events are also reported for miR844 which induces cleavage of its target *CDC3* transcript at 6, 12, 21, 52 nt upstream of the predicted target sites, resulting in the instability of the mRNA (Lee et al., 2015). It is also noteworthy that the target site in *WHY3* transcript is close to its PolyA tail (33 nt upstream, Figure S7B). Whether that accounts for the inhibition of translation efficiency is currently not clear. However, the widespread alternative lengthening of 3’ UTRs in most protein-coding genes and long non-coding RNAs are known to affect the functional stability, localization and translation efficiency of the RNAs. (Elkon et al., 2013; Chen et al., 2018). In fact, *WHY3* transcript exists in two 3’UTR by alternative polyadenylation (APA): the short one is 242 nt without the miR840 targeting region, while the long one is 563 nt in length (Figure S7B). Further clarification of the regulatory role of *WHY3* APA may be needed.

Target prediction using psRNATarget website with low stringency resulted in 121 and 128 candidate transcripts for the 22 nt miR840 and the 21 nt miR840*, respectively (Supplementary data set1). However, these target genes were not verified in the present study, and not reported in other publications concerning miR840 (Rajagopalan et al., 2006; Nodine and Bartel, 2010).

So far, miR840 is found only in Arabidopsis and close relatives, thus it appears to be an evolutionary young microRNA (Rajagopalan et al., 2006). The origin of miR840 may be of general interest in future studies (Cui et al., 2017), owing also to its special target configuration.

### The miR840-PPR/WHY3 module functions in plant senescence

During normal growth in Arabidopsis, accumulation of miR840 is found to associated with the development of senescence symptoms (Figure 2). Whereas knocking down miR840 could delay plant senescence, overexpression of miR840 enhances the senescent phenotype (Figure 3; Figure 4). At the molecular level, miR840 targets a convergent gene pair *PPR* and *WHY3* for either transcript degradation or translational repression, respectively and thereby reprograms many senescence-associated downstream genes (SAGs) (Figure S5; Figure S6). These include developmental signal-related *SAGs* like *WRKY53* (Miao and Zentgraf, 2007), *SAG12* (He and Gan, 2002), *SIRK* (Robatzek and Somssich, 2002), *SPO11-1, 2, 3* (Hartung et al., 2007) and *RAD52* (Samach et al., 2011), as well as environmental stress-induced *SAGs* such as *pyruvate decarboxylase* (Kursteiner et al., 2003), *glutaminesynthase* (Li et al., 2006), *CAMTA3* (Nie et al., 2012), *AHK3* (Kim et al., 2006), *RLP27* (Wang et al., 2008), *COR78* (Yang et al., 2011b), *WRKY70* (Besseau et al., 2012), *SAG101* (Feys et al., 2005), *WRKY33* (Birkenbihl et al., 2012), *EIN2* (Kim et al., 2009), *PDFs* (Liu et al., 2007) and *PR1* (Uknes et al., 1992; Epple et al., 1997; Thomma et al., 2002). Some of these downstream genes are involved in the response to wounding, jasmonic acid, fungus, water deprivation, cold and acid stresses (Alonso et al., 1999; Kim et al., 2006; Kim et al., 2009; Yang et al., 2011a; Thomas, 2013), implying that the miR840-induced early senescence may affect biotic and abiotic stress responses.

miR840-mediated senescence onset requires joint repression of both *WHY3* and *PPR*, due to the fact that single mutation of either *WHY3* or *PPR* is not sufficient to mimick the miR840 overexpression phenotype. However, the double mutant *kdwhy3 appr* displays a similar phenotype as the miR840 overexpressing plants, and that only co-expression of both in this mutant background can rescue the early senescent phenotype (Figure 8A-B). Consistently, several SAGs were found to share similarly regulated expression pattern among *kdwhy3*, *appr*, *kdwhy3 appr* and miR840 overexpressor mutants (Figure S5 and Figure S9).

Interestingly, both *PPR* and *WHY3* were not known for their involvement in plant senescence in previous studies, possibly owing to the weak phenotype of respective single mutants. *PPR* belongs to the pentatricopeptide repeat superfamily which encodes ~ 450 proteins in Arabidopsis. PPRs are RNA-binding proteins and are found in complexes of organelle mRNA-editing machineries in plants, which play roles in the regulation of photosynthesis, respiration, as well as in plant development and environmental responses (Barkan and Small, 2014). On the other hand, *WHY3* is a member of the three-gene family of WHIRLY single-stranded DNA binding proteins in Arabidopsis (Cappadocia et al., 2013) and is believed to localize dually in nucleus or plastid and mitochondria (Marechal et al., 2009; Jiang et al., 2017; Golin et al., 2020). Its closest homolog *WHY1* represses the senescence marker gene *WRKY53* and coordinates leaf senescence in a developmental stage-dependent manner (Miao et al., 2013; Huang et al., 2017; Ren et al., 2017; Huang et al., 2018; Lin et al., 2019).

Taken together, this work adds new regulatory aspects in plant development and onset of senescence depending on the evolutionary young miR840. Its ability to accelerate plant senescence upon its accumulation depends mainly on joint repression of the neighboring convergent genes *PPR* and *WHY3* by targeting their overlapping 3’ UTRs (Figure 9).

## Materials and Methods

### Plant materials and growth conditions

Plants of *Arabidopsis thaliana* (ecotype Columbia) were grown in a growth chamber under long (16 h light/8 h dark) or short (8 h light/16 h dark) illumination condition as described before (Miao et al., 2013). Phenotyping was assessed under the same illumination condition in all cases as indicated appropriately in the Results section.

T-DNA insertion lines *SAIL_232_F08* and *SALK_038777* for *MIR840 (AT2G02741), Salk_005345C* for *WHY3 (AT2G02740)* were ordered from NASC (http://arabidopsis.info/BasicForm) and confirmed by genotyping PCR with the primers suggested by the T-DNA Express Tool and by qRT-PCR with gene specific primers (Table S2). The homozygote *dcl1,* \ *dcl2, dcl3, dcl4, dcl4-2t* and *dcl2dcl4* mutants were kindly provided by other sientists. Plasmids for *Overexpression-PPR (35S::PPR)* and *overexpression-PPR-3’UTR (35S::PPR-UTR)* were created by inserting the *PPR* coding sequence alone or with its *3’UTR* into the destination vector pB2GW7 by GATEWAY cloning technology. Transgenic lines *appr/*WT and *kdwhy3 appr* were created by transforming WT and *kdwhy3* plants with the *antisense-PPR* (*appr*) plasmid constructed on pB2GW7 by Gateway-cloning the complementary *PPR* CDS (Figure 8A). Transgenics were selected by spraying seedlings with 0.1% (w/v) glufosinate ammonium and confirmed by semi-qPCR and qRT-PCR, and the T3 homozygous plants were used for phenotype observations. The *pri-miR840 OE* (Figure 4A) transgenic plants were generated by transformation of WT plants using pCBIM (Ren et al., 2012) harboring the 226 bp *pri-miR840* sequence (Figure 1A).

The tandem mimicry miR840 inhibitory vector was constructed in such a way that a nine-tandem-repeated miR840 22-nt sequence together with insertion of three extra nucleotides in the 10^th^ and 11^th^ position to form an unpaired inhibitory loop after transcription was first assembled (Figure 4A; Figure 4SA) and inserted reversely into the MCS site of the pCBIM vector, as described in detail previously (Jiang et al., 2014). The resulting plasmid and the pCBIM empty vector were used to generate the *9x miR840am* and the control transgenic plants on the WT background, respectively. The T1 seedlings were screened on plates containing 50 μg/ml hygromycin and the positives were checked with RNA gel blot (Figure 4B). Transgenic plants of *9x miR840am* showing an expected interfering RNA species (marked with a red arrow in Figure 4B) were selected for further experiments (Figure 5h).

### RNA isolation, northern blot, real-time quantitative PCR (qRT-PCR) and stem-loop qPCR

Total RNA was extracted using Trizol reagent (Invitrogen). Twenty micrograms of total RNA were used for long or small RNA isolation as described before with minor modification (Miao et al., 2004). Briefly, high molecular weight RNA was selectively precipitated from the total RNA by addition of one volume of 20% PEG with 1M NaCl (Llave et al., 2002). The supernatants enriched with low molecular weight RNA (80-100 μg) was then separated on 17% denaturing polyacrylamide gels and electro-blotted to a Hybond-N^+^ membrane for small RNA detection. Probes for miR840, siRNA1003 and U6 were synthetic oligonucleotides complementary to their sequences and end-labeled with ^32^P-ATP using T4 kinase (Fermentas). Probes for full-length *PPR* or *WHY3* CDS were ^32^P random-primer-labeled complementary DNA. Unincorporated nucleotides were removed using G-25 spin columns (Amersham) according to the manufacturer’s instruction. Blotting conditions were previously described (Miao et al., 2004). Membranes were exposed to X-ray films and the ethidium bromide stained rRNA and tRNA was used as loading amount control.

For qRT-PCR, first-strand cDNA was synthesized and detected using a RevertAid First Strand cDNA Synthesis kit (Thermo Fisher Scientific) and SYBR Green PCR Master Mix (Invitrogen). Quantification of expression were based on reference genes *ACTIN 2* or *PP2A* when developmental stages were compared. Gene-specific primers for *MIR840*, *PPR* and *WHY3*, as well as the fifty selected downstream genes were list in Figure S5A, and their specificity and amplification efficiency were confirmed by examining the melting curves.

For stem-loop qPCR analysis of microRNA, reverse transcription was done with miRNA first strand cDNA synthesis kit (Vazyme biotech, MR101-02) according to the manufacturer’s instruction. The specific stem-loop primer was designed by the online miRNA primer design software provided by the manufacturer, such that the RT primer for miR840 and U6-29 contained with their 5’ 6 nt annealing to the respective mature microRNA 3’ 6 nt, and the qPCR primers consisted with a universal reverse primer together with the microRNA-specific primers (Kramer, 2011). The primer sequences are given in Table S3.

### 5’RLM-RACE and sequencing

Using T4 RNA ligase 1 (NEB, M0437M), total RNA was isolated from 10-week-old or senescent rosettes of WT plants were ligated to a specific oligo adaptor at their 5’terminal, subsequently converted to first-strand cDNAs with a RevertAid first strand cDNA synthesis kit (Thermo Scientific™, K1622) following the manufacturer’s instruction. The detail procedure was described in supplementary method and Figure S6A. The RNA oligo and primers used in 5’RLM-RACE are listed in Table S4.

### GUS reporter assay for miR840-guided gene repression

A cloning plasmid pGWB433-MCS-GUS was constructed (Supplementary Method). This empty vector was assigned as reporter plasmid zero R(0) in Figure 5A. Oligonucleotides with an added ATG reminiscent of the 78 bp in the *WHY3 3’UTR* region overlapping with *PPR*, containing either the original (CACT) or mutated (GTGT) target sites of *WHY3,* were synthesized and inserted into pGWB433-MCS-GUS in frame with the *GUS* gene to create another 2 reporter plasmids R(W3UTR) and R(W3UTRm), respectively. Similarly, the reverse complementary sequence of the same 78 bp region was used to generated PPR 3UTR-GUS reporter plasmids R(P3UTR) and R(P3UTRm) containing the original (TGCG) and mutated (ACTA) target sites, respectively (Figure 5A and Figure S7). For construction of the effector plasmid, full-length *pri-miR840* sequence was cloned into the MCS of the vector pCBIM (Ren et al., 2012). All constructs were verified by sequencing.

*Agrobacterium tumefaciens* strain GV3101 harboring individual plasmid was grown overnight in liquid LB medium containing 20 μg/mL Acetosyringone (AS) and a diluted culture with 100 μg/mL AS reaching an OD_600_ 2.0 was harvested for leaf infiltration of one-month-old *N. benthamiana* plants after washing and resuspension with MES-KOH buffer (pH 5.7) at a concentration of OD_600_ 0.4. For co-infiltration, equal amount of Agrobacterium suspensions was used. Before infiltration of the 3rd leaves of *N. benthamiana,* the Agrobacterium suspensions was kept on bench for 3 hrs. At 3 days post infiltration, leaf discs were sampled for GUS staining and qRT-PCR verification. The plasmid-based *HptII* was used as reference gene for expression quantification. The primers used in this experiment are listed in Table S4.

### Inducible luciferase-fusion assay

To construct the WHY3-LUC fusion, a pENTR/TOPO-WHY3-LUC vector with the 560 bp *WHY3* 3’UTR sequence (UTR), the mutated version of the 3’UTR (UTRm) or without the 560 bp WHY3 3’UTR sequence were created. The mutated version of the 3’UTR *(UTRm)* was produced by incorporating substitute nucleotides in the reverse PCR primer. The LUC-UTR or LUC-UTRm cassette, was then isolated by digesting with *Not*I and *Eco*RI and sub-cloned into a previously constructed gateway entry vector harboring the full-length CDS of *WHY3*, to yield pENTR/TOPO-WHY3-LUC-UTR or pENTR/TOPO-WHY3-LUC-UTRm such that the *LUC* CDS was downstream in frame with the *WHY3* CDS linked by 21 bp sequence including the *Not*I site (Figure 7B). Finally, three binary vectors were generated by LR reaction using pMCD7 (Curtis and Grossniklaus, 2003) and the above entry vectors. All constructs were subjected to sequencing verification.

Transgenic seedlings were selected by spraying with 0.1% (w/v) glufosinate ammonium and identified by semi- and quantitative real-time PCR. Luciferase activity was determined according to the instruction manual of the reporter assay system (Promega, USA) with modifications. Briefly, 100 mg leaf discs were harvested and frozen in liquid nitrogen. After grinding, 100 μL 1x passive lysis buffer was added and mixed vigorously. The samples were incubated for 1 h on ice followed by centrifugation for 20 min at 13,000 x g. The resulting supernatant was diluted 1:5, 1:10, 1:20, 1:40 and put on a 96-well plate. After subsequently adding LARII, luminescence was measured in a Flexstation 3 Microplate Reader (Molecular Devices, USA). Total protein in the supernatants were determined by the Bradford method. Experiment was repeated at least three times.

### Protein extraction and immunodetection

For total soluble protein extraction, 200 mg fresh leaf materials were batch frozen in liquid nitrogen, ground into powder, resuspended in 100 μL of extraction buffer (100 mM Tris, pH7.2, 10% sucrose, 5 mM MgCl_2_, 5 mM EGTA, protease inhibitor) and centrifuged at 15,000 x g for 10 min. The supernatants were used for western blot analysis. Proteins were separated on 10% acrylamide gel and transferred to nitrocellulose membranes by semi-dry blotting. The membranes were blocked for 1 h at room temperature in TBS buffer containing 5% (w/v) non-fat dry milk powder, then incubated either with anti-WHY3 peptide antibody (provided by Prof. Dr. Karin Krupinska, University of Kiel) or with antibody-free PBS solution for 1 h, respectively. Blots were washed in TBST buffer for 10 min (3 times) before incubation with secondary antibody conjugated with a peroxidase. The blots were washed with TBST for 10 min (3 times) and then detected by chemiluminescence.

### Measurements of chlorophyll and carotenoid content, chlorophyll fluorescence and membrane ion leakage

The 7^th^ true leaf of a rosette was used for chlorophyll extraction. Chlorophyll a, b, total chlorophyll and total carotenoids contents were calculated according to a previous reported method (Wellburn, 1994). At least 15 plants were determined to calculate the representative mean of the biological sample. In some cases, chlorophyll contents were also detected by using the DUALEX^®^ SCIENTIFIC+ portable plant polyphenol-chlorophyll meter (Force-A, France), and the results were presented in unit μg/cm^2^. Chlorophyll fluorescence was measured from 5- to 13-week-old plants (grown under 8 h light) as described previously (Miao et al., 2013).

The No.7 true leaf in 5- to 13-week-old rosettes was used for measurement of membrane ion leakage as described previously (Miao et al, 2007).

### Statistical analysis

Mean values and standard deviations (SD) were calculated in Microsoft Office Excel 2019. Statistical significance among various comparisons were analyzed by one-way ANOVA or pair-wide multiple t-tests using the software Origin 7.5 (OriginLab Corporation, USA). Two asterisks indicate extremely significant differences when *p*-value ≤ 0.01, while one asterisk indicates significant differences with a *p*-value ≤ 0.05.

## Acknowledgements

We express our gratitude to Prof. Dr. Ulrike Zentgraf, University of Tuebingen, and Prof. Dr. Karin Krupinska, University of Kiel, for kindly providing a peptide antibody against WHY3. and Prof. Dr. Jian-Kang Zhu (University of California, USA) for providing the *dcl1* mutants and Prof. Dr. Yijun Qi (Tsinhua University) for the *dcl2, dcl3, dcl4* and *dcl2/dcl4* mutants. Prof. Dr. Binghua Wu is appreciative for his critical comments and helpful suggestions. This work was supported by the National Natural Science Foundation of China (grant no. 31770318, 31470383, and 31400260), the German Research Foundation (DFG: MI1392/1-1), and the financial support from China Scholarship Council to Y.R. and international exchanging program of Fujian Agriculture and Forestry University (KXB16009A).

## Author contributions

Y.M. and Y.R. designed the project. Y.R., Y.M., W.W., and W.L. performed the experiments, collected data, and analyzed the results. Y.R. and Y.M. wrote the manuscript. D.C. and S.D. contributed to miRNA experimental discussions and manuscript correction.

## Data availability

The authors declare that all data supporting the findings of this study are available within the manuscript and its supporting information is available from the corresponding author upon request.

**Supplementary materials (figure legends, captions of S. Tables and data set)**

**Figure S1 Expression quantification of miR840, *PPR* and *WHY3* in leaf position of a 13-week-old rosette**. The value was displayed as a mean ± SD from three biological replicates.

**Figure S2 Differential senescent area in a single leaf and gene expression analysis.** * *p* < 0.05 ** *p* < 0.01 (n =3).

**Figure S3 Distribution of *cis*-acting elements in the −1000 bp region of *MIR840* promoter.** The location of each *cis*-acting element is colored and labeled below the sequences. The T-DNA insertion sites of the gain-of-function *(SAIL_232_F08)* and loss-of-function *(SALK_038777)* mutants were indicated by arrow line.

**Figure S4 Illustration for mimicry miR840 inhibition, *WHY3* knockdown T-DNA mutant and the *WHY3* overexpression construct. (A)** A working model of the mimicry miR840. The reverse mimicry miR840 finds and anneals to the normal miR840 *in vivo*, leading to competitive inhibition the functions of the normal miR840. **(B)** Semi-qPCR to detect the expression level of the 9x tandem mimicry miR840 construct (*9x miR840am)* in T1 transgenic plants compared to WT control. Red arrow indicates the positive band in different transgenic line. **(C)** The T-DNA insertion site of the *kdwhy3* mutant. **(D)** The *overexpression WHY3* (*oeWHY3*) construct. *P35S*, *CaMV35S* promoter; *Tnos, NOS* terminator; *WHY3,* the full length *WHY3* CDS.

**Figure S5. Quantification of expression of 50 known senescence-associated genes in rosette of the T-DNA insertion lines *SAIL_232_F08* (with up-regulated miR840) and *SALK_038777* (with down-regulated miR840).** Related information of the selected genes with reference is listed in (A). Data represents fold-change over that of WT (n = 3 plants).

**Figure S6 Determination of *miR840*-guided cleavage sites in *PPR* and *WHY3* transcripts using 5’RLM-RACE cloning and sequencing. (A)** Qualitive control of RNA preparation, adaptor ligation (left panel) and reverse transcription reaction (RT) in the RLM-RACE system (middle and right). The known high abundantly expressed *UBQ13* (middle panel) and lowly expressed gene *SUC7* (right panel) were amplified from the cDNA preparations. **(B)** The *WHY3* and *PPR* RACE products were amplified by 2-round PCR using gene specific primer together with 5’RACE primer (lane 1) and with 5’nested primer (lane 2) subsequently. The *NAC1* transcript targeted by *miR164* serves as method verification.

**Figure S7 Nucleotide sequences of the *3’UTR* of *PPR* (A) and *WHY3* (B) showing the localization of the miR840-guided cleavage sites (blue and red arrow) and binding regions of miR840/miR840* (boxed).** The 78 nt sequences used for synthetic oligonucleotides in the GUS reporter assay are indicated by brackets below the sequences.

**Figure S8 Senescence-related phenotyping of the *knockdown why3 (kdwhy3)* and *antisense-PPR (appr)* transgenic lines. (A)-(C)** Semi-qPCR and qRT-PCR verification of the transcript level of *PPR* in the *appr* transgenic lines and of *WHY3* in the *kdwhy3* mutant. T3 homozygous plants were used. The value is displayed as a mean ± SD of three biological replicates. Significant differences were detected by one-way ANOVA test using Origin 7.5 software. ** *P* < 0.01. **(D)** Comparison of senescent status of *appr*, *kdwhy3* and WT plants grown for 8 weeks in short illumination condition. Rosette leaves were arranged in the order from the oldest to the youngest. **(E)-(F)** Fv/Fm value and total chlorophyll content of the 7^th^ rosette leaf measured during development. The value is displayed as a mean ± SD of twelve independent measurements. **(G)** Ratio of categorized leaves in 8-week-old plants. The value represents mean ± SD of 12 independent measurements.

**Figure S9 Quantification of gene expression of selected senescence-associated genes in rosettes of *kdwhy 3appr*, *kdwhy3* and *appr* transgenic plants, shown as fold change over that in WT.** mean ± SD (n = 3 plants). The list of genes is shown in Figure S5.

**Supplementary data set 1. miR840 target prediction by psRNATarge date20200718.**

**Table S1. Segregation of senescence-like phenotypes in different progeny generations of SALK_038777 and SAIL_232_F08 plants.**

**Table S2. Primers used for semi- or qRT-PCR analysis of gene expression in this research.**

**Table S3. Primers used for stem-loop qPCR of miR840 and related vector constructions.**

**Table S4. Primers used for 5’RLM-RACE reactions and GUS reporter assay.**

